# Impaired glymphatic clearance independently contributes to poor outcomes in Parkinson’s disease

**DOI:** 10.1101/2025.01.15.633106

**Authors:** Angeliki Zarkali, George Thomas, Ross Paterson, Naomi Hannaway, Ivelina Dobreva, Amanda J Heslegrave, Elena Veleva, Henrik Zetterberg, Rimona S Weil

## Abstract

**Background:** Impaired glymphatic clearance may contribute to pathological accumulations in Parkinson’s (PD), but how it interacts with other processes causing dementia and poor outcomes remains unclear.

**Objectives:** Clarify how glymphatic clearance impacts cognition in PD and its interaction with established imaging markers.

**Methods:** We used diffusion tensor image analysis along the perivascular space (DTI-ALPS) as an indirect marker of glymphatic clearance in 98 PD patients (31 PD-poor outcomes: dementia, mild cognitive impairment, frailty or death within 3-year follow-up; 67 PD-good outcomes) and 28 controls. We assessed DTI-ALPS relationship to cognition, white matter (fibre cross-section), cortical thickness, iron accumulation (quantitative susceptibility mapping (QSM)), and plasma markers (phosphorylated tau-181 (p-tau181 and neurofilament light (NFL)) cross-sectionally and longitudinally.

**Results:** DTI-ALPS was lower in PD-poor outcomes compared to PD-good outcomes and controls (p=0.005) with further longitudinal reductions only in PD-poor outcomes (group*time interaction: β=-0.013, p=0.021). Lower DTI-ALPS was associated with lower fibre cross-section in P, at baseline and longitudinally but with different spatial distribution from white matter changes relating to PD cognition. There was no correlation between baseline DTI-ALPS and plasma ptau-181 (p=0.642), NFL (p=0.448) or baseline cortical thickness. Lower DTI-ALPS was associated with accelerated cortical thinning within left precentral gyrus and changes in brain iron distribution.

**Conclusions:** PD patients who develop poor outcomes show impaired glymphatic clearance at baseline that worsened longitudinally. DTI-ALPS correlated with white matter integrity and brain iron accumulation. However, both showed different spatial distribution than that seen in PD dementia; suggesting impaired glymphatic clearance contributes to cognitive decline in a distinct manner.

## Introduction

Cognitive impairment is common in Parkinson’s disease (PD), with dementia developing in nearly half of PD patients within 10 years^1^ with high personal, societal and financial burden^2,3^. PD dementia is multifactorial: outside of dopaminergic degeneration and alpha-synuclein accumulation, other processes including white matter degeneration^4,5^, iron accumulation^6^, beta-amyloid, tau co-pathology^7^ and concurrent cerebrovascular disease^8^ contribute to the development and progression of cognitive decline.

Additionally, recent studies have implicated glymphatic system dysfunction in the pathophysiology of PD and other neurodegenerative disorders^9^. The glymphatic system, first proposed by Iliff et al^10^, is a network of perivascular channels that facilitates flow of cerebrospinal fluid (CSF) through the brain and plays a key role in clearance of soluble proteins and waste through astrocytic aquaporin-4 channels^10,11^. In mouse models, monomeric alpha-synuclein propagates to remote brain regions via the glymphatic system^12^, and suppression of glymphatic system activity via aquaporin-4 gene deletion or acetazolamide administration leads to increased alpha-synuclein accumulation^13^. Similarly, impaired glymphatic system clearance is seen in mouse models of tauopathies^14^ and leads to beta-amyloid accumulation and cognitive deficits in mice^15^.

Techniques to assess the glymphatic system in-vivo have recently been developed, however most are invasive, requiring injection of tracers and sequential imaging^16^. Amongst non-invasive methods is diffusion tensor image analysis along the perivascular space (DTI-ALPS). This is thought to reflect glymphatic clearance by measuring diffusivity of water molecules in the perivascular space at the level of the body of the lateral ventricle^17^. Although an indirect measure from a single brain region aimed to reflect whole-brain clearance, DTI-ALPS strongly correlates with gold-standard measures of glymphatic function^18^ and conventional MRI markers of cerebrovascular disease^19^. Additionally, DTI-ALPS has good cross-vendor, inter-rater and test-retest reliability^20^. It recently showed positive correlations with sleep quality, younger age and better cognition in health^21^, in accordance with what would be expected from a metric of glymphatic clearance.

Several studies have shown reduced DTI-ALPS index in PD^22^, including in idiopathic rapid eye movement (REM) sleep behaviour disorder, reflecting prodromal synucleinopathy^23^. DTI-ALPS is also inversely correlated with motor symptoms and disease severity^24,25^ and linked to faster motor and cognitive decline in PD^25^. How and when impaired glymphatic clearance impacts other processes contributing to PD dementia is not clear, neither is the interrelationship between DTI-ALPS and other imaging metrics that change with cognitive impairment in PD^26^.

However, there is evidence that these may interact. In Alzheimer’s, where DTI-ALPS is also reduced and correlates with cognition^17,27–29^, DTI-ALPS is linked to beta-amyloid^29^ and to tau PET burden^28,29^. It also correlates with grey matter atrophy which may even mediate its effect on cognition^28–30^. In healthy older adults, higher DTI-ALPS index correlates with higher grey matter volume^31^, and relates to iron deposition within the basal ganglia^32^, assessed using quantitative susceptibility mapping (QSM). In PD, co-pathologies are particularly prevalent^33^ with high heterogeneity in cognitive presentation and underlying pathology. Single biomarkers have been unsuccessful in risk stratification and disease progression^26^. A more fruitful approach is to elucidate the complex interactions between pathological protein accumulation, tissue changes such iron build-up, small vessel disease, and glymphatic clearance, and how these contribute to cognitive impairment.

Here we used DTI-ALPS as an indirect measure of glymphatic clearance, and assessed its ability to predict poor outcomes (development of dementia, mild cognitive impairment (MCI), frailty or death over 3-year follow-up) in 98 PD patients and 28 controls. We assessed how glymphatic function interacts with other processes linked to poor outcomes in PD both cross-sectionally at baseline and longitudinally over 3 years. Specifically, we examined DTI-ALPS relationship with: 1) white matter macrostructural integrity, assessed using whole-brain fibre cross-section analysis and plasma neurofilament light (NFL); 2) grey matter atrophy, assessed using cortical thickness; 3) iron accumulation, using QSM; and 4) beta-amyloid and tau accumulation, assessed using plasma levels of phosphorylated tau at threonine 181 (p-tau181).

## Methods

### Participants

We included 98 PD patients and 28 controls from the Vision in Parkinson’s disease study, University College London; the study protocol has been previously detailed^34^. PD patients fulfilled Movement Disorders Society (MDS) clinical diagnostic criteria^35^. Participants were assessed at baseline, and after 18-(Session 2) and 36-months (Session 3). All participants provided written informed consent; the study was approved by the Queen Square Research Ethics Committee.

Clinical and neuropsychological assessments were performed whilst participants were receiving their usual medications to minimise attrition and discomfort. Neuropsychology assessments included two general measures of cognition, the Mini-Mental State Examination^36^ (MMSE) and Montreal Cognitive Assessment^37^ (MoCA), and two tests per cognitive domain^34^. A composite cognitive score was calculated as the averaged z-scores of each participant’s MoCA score plus one task per cognitive domain^34^. Motor function was assessed using MDS-UPDRS^38^ and Timed Up and Go test^39^ (TUG). Levodopa equivalent daily doses (LEDD) were calculated for PD participants^40^.

As in previous work^4,34^ we defined poor cognitive outcomes in PD if any of the following occurred during follow-up: death, frailty, dementia or mild cognitive impairment (PD-MCI) defined as persistent performance below 1.5 standard deviations in either 2 tests in one cognitive domain or 1 test in 2 cognitive domains^41^.

### MRI acquisition

MRI data for all sessions and participants were acquired on the same 3T Siemens Prisma scanner (Siemens) with a 64-channel head coil and parameters: *Diffusion weighted imaging (DWI):* b0 (AP- and PA-directions), b=50s/mm^2^ 17 directions, b=300s/mm^2^ 8 directions, b=1000s/mm^2^ 64 directions, b=2000s/mm^2^ 64 directions, 2 mm^3^ isotropic voxels, TE=3260ms, TR=58ms, 72 slices, acceleration factor=2; *3D magnetization prepared rapid acquisition gradient echo (MPRAGE):* 1mm^3^ isotropic voxels, TE=3.34ms, TR= 2530 ms, flip angle=7°; *Susceptibility weighted imaging (SWI)*: 1m^3^ isotropic voxels, 3D flow-compensated spoiled-gradient-recalled echo sequence, flip angle=12°, TE=18ms, TR=25ms, receiver bandwidth=110Hz/pixel, matrix size=204×224×160.

### MRI quality control and pre-processing

For all modalities, raw volumes were visually inspected and excluded if artefact was present in over 15 volumes. Images passing quality assurance underwent preprocessing using established pipelines, as previously described^4,42^.

Briefly, for *<u>white matter imaging</u>*, DWI images were pre-processed using the Mrtrix3 pipeline including denoising^43^, Gibbs artefact removal^44^, eddy-current, motion^45^, and bias field correction^46^; images were up-sampled (1.3mm^3^) and intensity normalisation performed. Multi-shell 3-tissue constrained spherical deconvolution^47^ was performed and fibre-orientation distributions calculated per participant and registered to a group-averaged template^48^. Fibre cross-section was derived for each participant; this was chosen as it is the most sensitive metric in PD^4,49,50^ and reflects pathological protein accumulation and subsequent neurodegeneration, rather than cerebrovascular changes^51^.

For <u>*grey matter imaging*</u>, T1-weighted images were processed with the default cross-sectional parameters using FreeSurfer6.0, then the longitudinal pipeline^52^, manually correcting and reprocessing surface reconstructions where necessary.

For *<u>QSM</u>*, 3D-complex phase data were unwrapped from SWI images using rapid path-based minimum spanning tree algorithm (ROMEO)^53^ and brain masks calculated using BET^54^. Background field removal was performed using Laplacian boundary value^55^ and dipole inversion was performed using multi-scale dipole inversion^56^ to derive susceptibility maps.

### DTI-ALPS index calculation

DTI-ALPS was calculated in Mrtrix3 using custom scrips based on Taoka et al^17^. DTI-ALPS assumes that movement of interstitial fluid in the perivascular space at the level of the body of the lateral ventricle is dominant parallel to the medullary veins, which run perpendicular to the ventricular wall; this is the x-axis (right-left direction). The y-axis (rostral-caudal) direction contains the projection fibres, and z-axis (anterior-posterior direction) the association fibres, orthogonal to the direction of the perivascular space^17^. DTI-ALPS index is defined as:

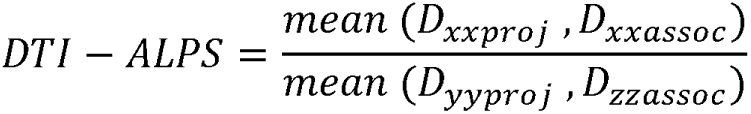

where D_xxproj_ is mean diffusivity in the area of projections fibers in the x-axis, D_xxassoc_ mean diffusivity in the area of association fibres in the x-axis, D_yyproj_ mean diffusivity in the area of projection fibres in the y-axis, and D_zzassoc_ mean diffusivity in the area of association fibres in the z-axis^17^.

We used an automated atlas-based approach to identify projection and association fibres based on a pipeline with good inter-rater, inter-scanner and test-retest reliability^20^. Projection fibres were defined using the superior corona radiata (SCR) in the John Hopkins (JHU) atlas (centre coordinates left (116,110,99), right (64,110,99)) and association fibres using the superior longitudinal fasciculus (SLF; left (128,110,99), right (51,110,99)).

The DTI-ALPS index was calculated separately for the left and right hemispheres and a mean index was calculated as the average of the two. As DTI-ALPS between hemispheres was highly intercorrelated (Spearman rho=0.819, p<0.001) we report the mean index, referred hereby as DTI-ALPS index. Lower DTI-ALPS index reflects lower diffusivity and worse glymphatic clearance.

### Plasma biomarkers

Plasma samples were collected at Session 2 and results were available for 87 participants for p-tau181 and 88 participants for NFL. 13 participants had plasma collected at Session 3 due to COVID-19 lockdown restrictions during Session 2 testing period. P-tau181 was measured using the Simoa p-tau181 Advantage Kit, and NFL using the Simoa Human Neurology 4-Plex A (N4PA) assay (Quanterix); analysts were blinded to clinical data. All measurements were above assay limit of detection. Quality control samples had mean intra-assay and inter-assay coefficient of variation of <10%.

### Statistical analysis

Group differences were assessed using ANOVA (post-hoc Dunn) for normally distributed and Kruskal Willis (post-hoc Tukey) for non-normally distributed variables. To assess the relationship of DTI-ALPS to clinical and other imaging metrics we used partial Spearman correlation cross-sectionally, and repeated measures ANOVA longitudinally correcting for age and sex.

To investigate the relationship *<u>between glymphatic function and white matter degeneration</u>* we performed whole-brain fixel-based analysis of fibre cross-section. Whole-brain probabilistic tractography was performed on the population template with 20 million streamlines, filtered to 2 million using spherical deconvolution-informed filtering of tractograms (SIFT)^57^. Connectivity-based fixel enhancement was performed on white matter fixels of the JHU atlas^58^ (5000 permutations, family-wise error (FWE)-corrected p-value q<0.05 and 10 voxels extent-based threshold). As DTI-ALPS is derived from DWI images, to minimise potential bias, we used the left DTI-ALPS index to correlate with right hemisphere fixels and the right DTI-ALPS index for left hemisphere fixels and report combined results. The following analyses were performed: 1) baseline fibre cross-section against baseline DTI-ALPS (baseline age, sex and total intracranial volume as covariates), and 2) follow-up fibre cross-section against baseline DTI-ALPS (baseline age, sex, total intracranial volume and time-between-scans as covariates).

To investigate the relationship *<u>between glymphatic function and cortical thickness</u>*, we used general linear models as implemented in Freesurfer. The following analyses were performed with same nuisance covariates as above: 1) baseline DTI-ALPS against baseline cortical thickness and 2) baseline DTI-ALPS against longitudinal cortical thickness. Significance maps were corrected for multiple comparisons using false discovery rate (FDR) combined over left and right hemispheres.

To relate *<u>DTI-ALPS to QSM signal</u>*, we first performed spatial standardisation of susceptibility maps using an average T1-weighted study template from included participants’ MPRAGE images^42^. Each participant’s bias-corrected magnitude image was registered to the respective MPRAGE (rigid and affine registration). QSM images were transformed to template space through high-order b-spline interpolation. Images were smoothed using a 3D

Gaussian kernel (3mm standard deviation) and smoothing compensated. Whole-brain QSM analysis was performed using permutation with threshold-free cluster enhancement (fslrandomise, 10,000 permutations, FWE-corrected q<0.05). The following analyses were performed with above nuisance covariates: 1) baseline susceptibility against baseline DTI-ALPS, and 2) follow-up susceptibility against baseline DTI-ALPS.

To explore the interplay between DTI-ALPS, white and grey matter degeneration, iron accumulation and poor cognitive outcomes in PD, we conducted mediation analyses to derive total, direct and indirect effects. These are detailed in *Supplementary Material*.

## Data sharing

Anonymised, group-level summary data and analysis code is available on GitHub (https://github.com/AngelikaZa/AngelikaZa-DTIALPSinPDcognitiveImpairment). Imaging and clinical data will be shared upon reasonable request to the corresponding author.

## Results

98 PD participants and 28 controls were included; demographics and clinical assessments are seen in *Table 1*.

**Table 1.** Baseline demographics and clinical assessments.

|  | Healthy Controls<br>n=28 | PD with good outcome<br>n=67 | PD with bad outcome<br>n=31 | p-value |
| --- | --- | --- | --- | --- |
| **Demographic characteristics** |  |  |  |  |
| Age | **65.7 (9.1)** | **62.4 (7.0)** | **68.5 (8.5)** | **p=0.004<sup>c</sup>** |
| Sex (F/M) | **15/13** | **38/29** | **8/23** | **p=0.014<sup>c</sup>** |
| Handedness (R/L) | 25/3 | 63/4 | 27/4 | p=0.516 |
| Years in Education | 17.9 (2.3) | 16.9 (2.6) | 17.5 (2.9) | p=0.386 |
| Total Intracranial Volume | 1393.7 (95.5) | 1459.7 (135.5) | 1468.9 (120.5) | p=0.036<sup>ns</sup> |
| **Clinical assessments** |  |  |  |  |
| MMSE | **29.2 (0.9)** | **29.1 (1.1)** | **28.4 (1.5)** | **p=0.004<sup>b,c</sup>** |
| MOCA | **28.9 (1.3)** | **28.5 (1.4)** | **26.3 (3.2)** | **p<0.001<sup>b,c</sup>** |
| Combined cognitive score | **0.07 (0.6)** | **- 0.01 (0.6)** | **-0.93 (0.9)** | **p<0.001<sup>b,c</sup>** |
| Disease duration | - | 4.1 (2.3) | 4.9 (3.4) | p=0.209 |
| Affected side at onset (R/L/BL) | - | 35/6/26 | 14/7/10 | p=0.082 |
| UPDRS total score | - | **41.4 (18.7)** | **55.6 (32.9)** | **p=0.007** |
| UPDRS motor score | - | **19.5 (9.5)** | **26.6 (18.8)** | **p=0.042** |
| LEDD | - | 451.8 (271.4) | 484.3 (231.7) | p=0.195 |
| Sleep (RBDSQ) | - | 4.1 (2.5) | 4.5 (2.5) | p=0.224 |
| **Image quality metrics** |  |  |  |  |
| Coefficient of joint variation* | 0.7 (0.2) | 0.7 (0.2) | 0.8 (0.2) | p=0.206 |
| Total signal to noise ratio* | 47.9 (12.1) | 47.4 (14.6) | 42.5 (15.2) | p=0.153 |
| Entropy focus criterion | 135.5 (28.0) | 131.1 (25.5) | 127.6 (24.4) | p=0.082 |

All results are shown as mean (standard deviation) unless otherwise specified.
PD: Parkinson's disease, F: Female, M: Male, R: Right, L: Left, BL: Bilateral; MMSE: Mini Mental State Examination; MOCA: Montreal Cognitive Assessment; Combined Cognitive Score: calculated as the mean of the z scores of 2 cognitive scores per cognitive domain (z-scored against control performance at baseline); UPDRS: Unified Parkinson's Disease Rating Scale, LEDD: Total equivalent Levodopa Dose, RBDSQ: REM Sleep Behaviour Disorder Questionnaire.
<sup>a</sup>post-hoc significant difference between HC and PD good outcome; <sup>b</sup>post-hoc significant difference between HC and PD poor outcome; <sup>c</sup>post-hoc significant difference between PD good and PD poor outcome (also in bold). \*lower values are better.

### Lower baseline DTI-ALPS index is associated with poor outcomes in PD

PD poor outcomes showed lower DTI-ALPS index compared to PD good outcomes and controls (PD poor outcomes mean±SD= 1.08±0.16 vs PD good outcomes 1.19±0.16 vs controls 1.18±0.20, p=0.005)(*Figure 1B*). PD poor outcomes showed further longitudinal reductions (group*time interaction β=-0.057, p<0.001) within 18-month follow-up (*Figure 1B*). Baseline DTI-ALPS index was correlated with combined cognitive scores at baseline (rho=0.226, p=0.027) (*Figure 1C*) and Session 3 (rho=0.25, p=0.014)(*Figure 1D*) but not longitudinal change in combined cognitive scores (p=0.088).

**Figure 1.**
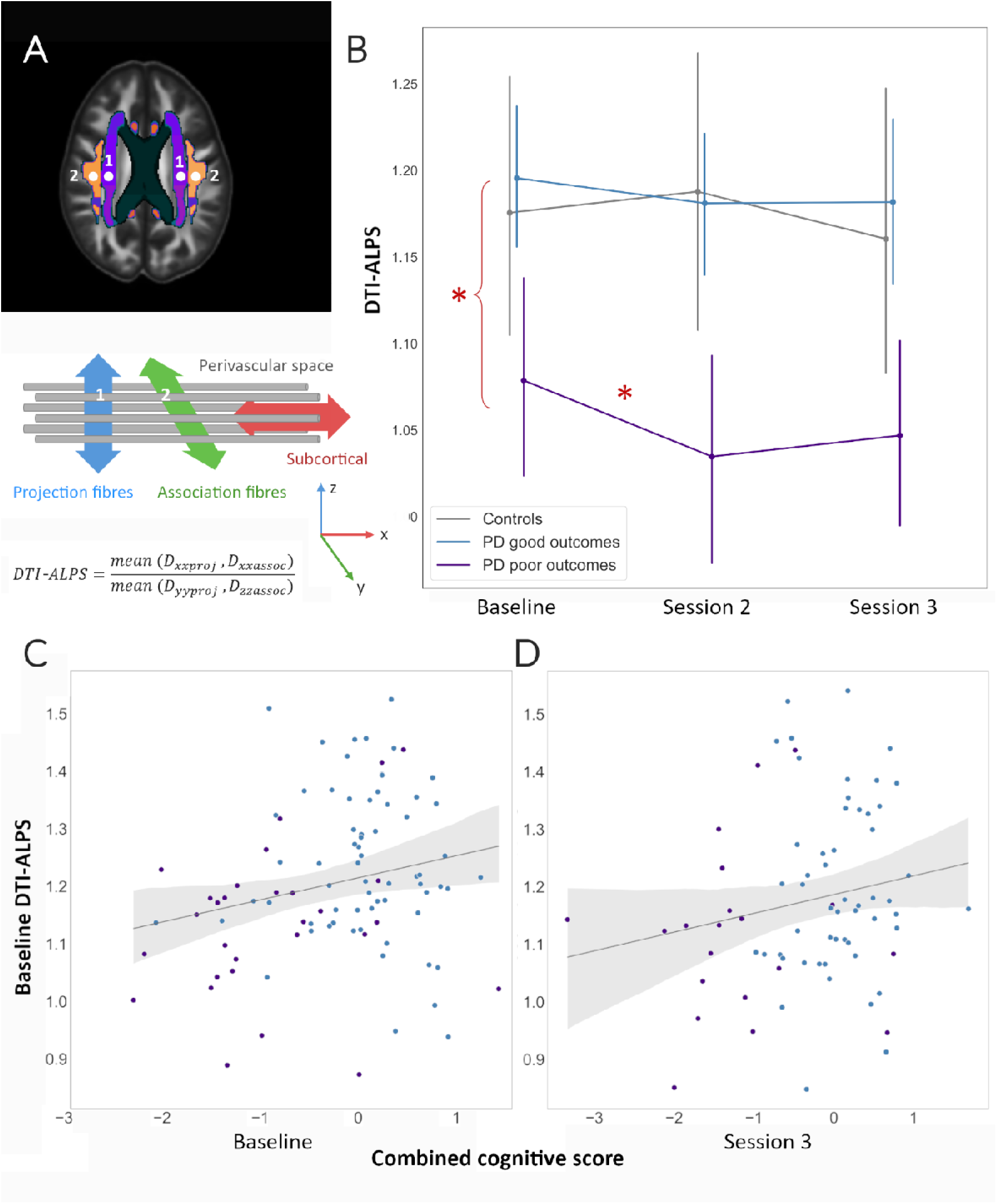
DTI-ALPS index is associated with poor cognitive outcomes in Parkinson’s disease. **A. Calculation of DTI-ALPS.** DTI-ALPS is based on the assumption that diffusion of fluid in the perivascular space at the level of the body of the lateral ventricle is dominant parallel to the medullary veins which run along the x-axis. Lower values indicate impaired glymphatic clearance. The diagram shows the positional relationship of the perivascular space (x-axis), projection fibres (z-axis) and association fibres (y-axis). Their orthogonal relationship makes it possible to evaluate the diffusion component along the medullary vessels by eliminating the influence of the fibre component using the formula:

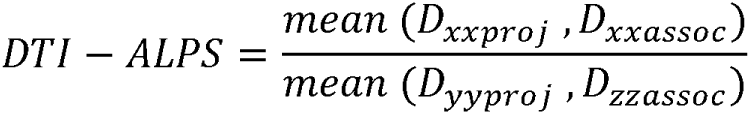 We identified projection (1) and association (2) fibres at the level of lateral ventricle body as 5mm spheres within the superior corona radiata (SCR) and the superior longitudinal fasciculus (SLF) of the JHU-ICBM-DTI-81-white-matter labelled Atlas. ROI centre coordinates were SCR: left (116,110,99) right (64,110,99) and SLF: left (129,110,99) and right (51,110,99) in the JHU-ICBM-FA template. **B. DTI-ALPS index is lower in patients with Parkinson’s who progressed to poor outcomes.** Patients with Parkinson’s who had poor cognitive outcomes had lower DTI-ALPS index (indicating impaired glymphatic clearance) at baseline (p=0.005) and showed additional longitudinal reductions (group*time interaction β -0.057, p<0.001) within 18 months follow up. **C. Baseline DTI-ALPS index was correlated with cognition at baseline**. Baseline DTI-ALPS index was correlated with combined cognitive scores (z-scores of MOCA plus one test per cognitive domain) at baseline in patients with Parkinson’s disease (rho=0.226, p=0.027). **D. Baseline DTI-ALPS index was correlated with cognition at 3-year follow-up.** Baseline DTI-ALPS index was correlated with combined cognitive scores at Session 3 (rho=0.25, p=0.014). However, baseline DTI-ALPS index was not correlated with longitudinal change in combined cognitive scores (p=0.088). DTI-ALPS: diffusion tensor image analysis along the perivascular space; PD: Parkinson’s disease; ROI: region of interest; SCR: superior corona radiata; SLF: superior longitudinal fasciculus.

Having confirmed the association between DTI-ALPS index and development of poor cognitive outcomes in PD, we aimed to assess how impaired glymphatic clearance (reflected by lower DTI-ALPS) interacts with processes known to correlate with cognitive impairment; specifically loss of white matter macrostructural integrity (assessed using fibre cross-section and plasma NFL), iron accumulation (assessed using QSM), amyloid co-pathology (assessed using plasma p-tau181) and grey matter atrophy (assessed using cortical thickness).

### Lower DTI-ALPS index is associated with changes in white matter macrostructure but with a different spatial distribution to changes linked to cognitive impairment in PD

First, we assessed whether DTI-ALPS index in PD was associated with loss of macrostructural integrity (fibre cross-section) at baseline (n=98) and after 3-years (Session 3, n=64), using whole-brain fixel-based analysis corrected for age, sex and intracranial volume (and time-between-scans for Session 3). Lower DTI-ALPS index was associated with reduced fibre cross-section within the right optic radiation, bilateral anterior corona radiata and left arcuate fasciculus at baseline (up to 33.6% reduction, FWE-corrected q=0.007). Lower DTI-ALPS was associated with reduced fibre cross-section within the left anterior and posterior corona radiata at Session 3 (40.0% reduction, q=0.006). White matter changes associated with DTI-ALPS are seen in *Figure 2*; these were spatially distinct from previously described changes linked to cognitive impairment in PD^4,49^ and independent from DTI-ALPS in mediation analyses (*Supplementary Material S1)*.

**Figure 2.**
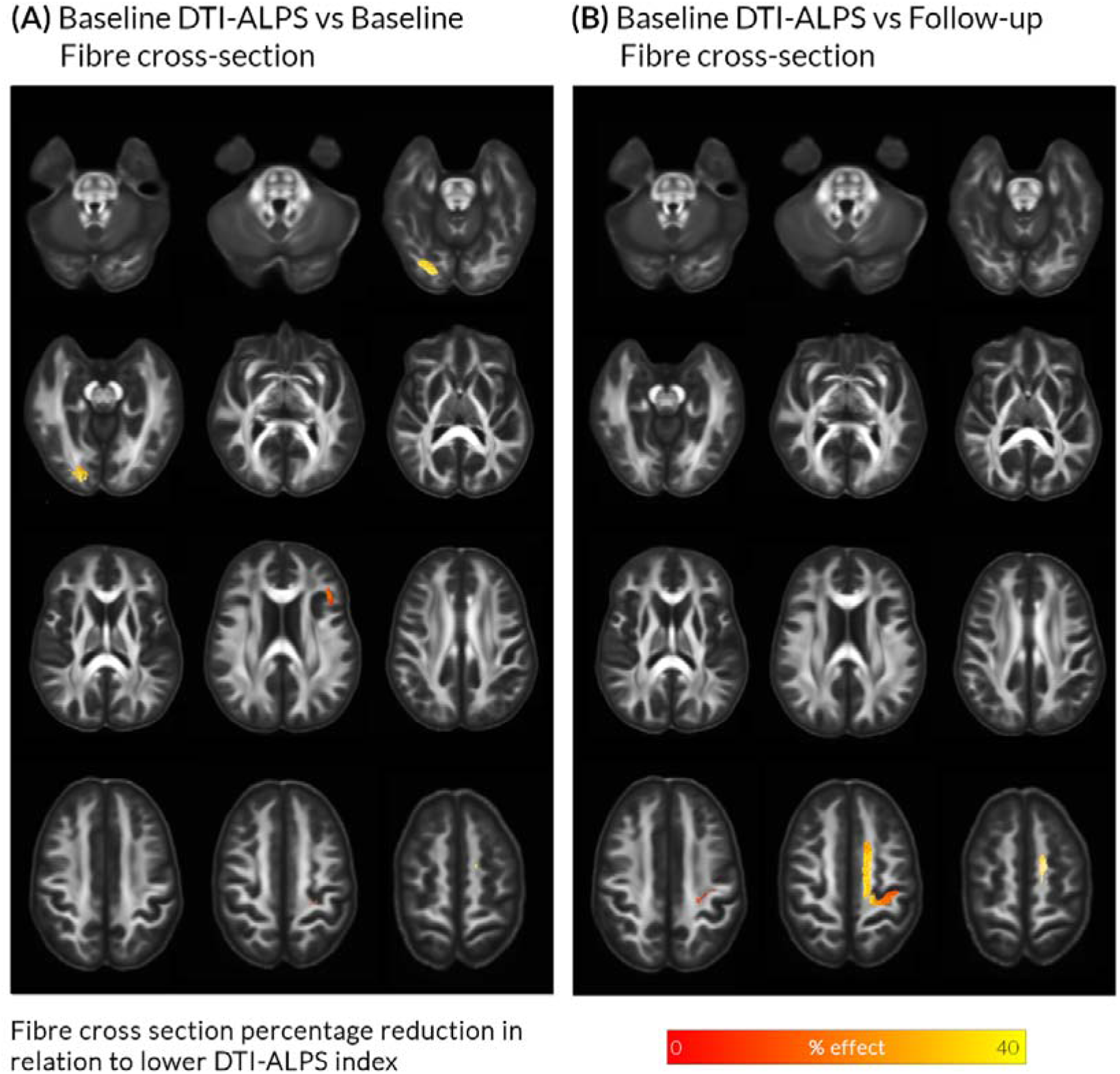
Lower DTI-ALPS index, indicating impaired glymphatic clearance, is associated with white matter macrostructural changes in Parkinson’s disease. **A. Baseline reductions in fibre cross-section in whole brain analysis.** In patients with Parkinson’s disease (n=98), lower baseline DTI-ALPS index was associated with reductions in baseline fibre cross-section, in occipital regions, adjusted for age, sex and intracranial volume. **B. Reductions in fibre cross-section after 3 year follow-up (Session 3).** In patients with Parkinson’s disease (n=64), lower baseline DTI-ALPS index was associated with reductions in fibre cross-section in superior frontal regions, at Session 3, adjusted for age, sex, total intracranial volume and time between scans. Effect sizes in both cases are shown as percentages and presented as streamlines in template space. Only family-wise-error (FWE) corrected results are displayed (FWE-corrected p<0.05). DTI-ALPS: diffusion tensor image analysis along the perivascular space.

Baseline DTI-ALPS index was not associated with plasma NFL in PD (n=82) (β=-0.351, p=0.448). There was also no significant correlation between DTI-ALPS and p-tau181 (n=88, β =-0.266, p=0=0.642).

### DTI-ALPS index is associated with QSM signal, reflecting brain iron re-distribution

We assessed the relationship between DTI-ALPS index and whole-brain QSM signal in PD at baseline (n=98) and Session 3 (n=59). For baseline QSM, lower DTI-ALPS index was associated with higher QSM signal within bilateral cerebellar peduncles, medial temporal, posterior parietal and frontal lobes, and lower QSM signal within the and bilateral occipital regions (*Figure 3*). Lower baseline DTI-ALPS was associated with higher QSM signal in right cingulate and posterior parietal lobe after 3-years follow-up (*Figure 3*). The spatial distribution of QSM signal change was distinct from previously described changes associated with worse cognition in PD^6,42^ and not mediated by DTI-ALPS (*Supplementary Material S1*).

**Figure 3.**
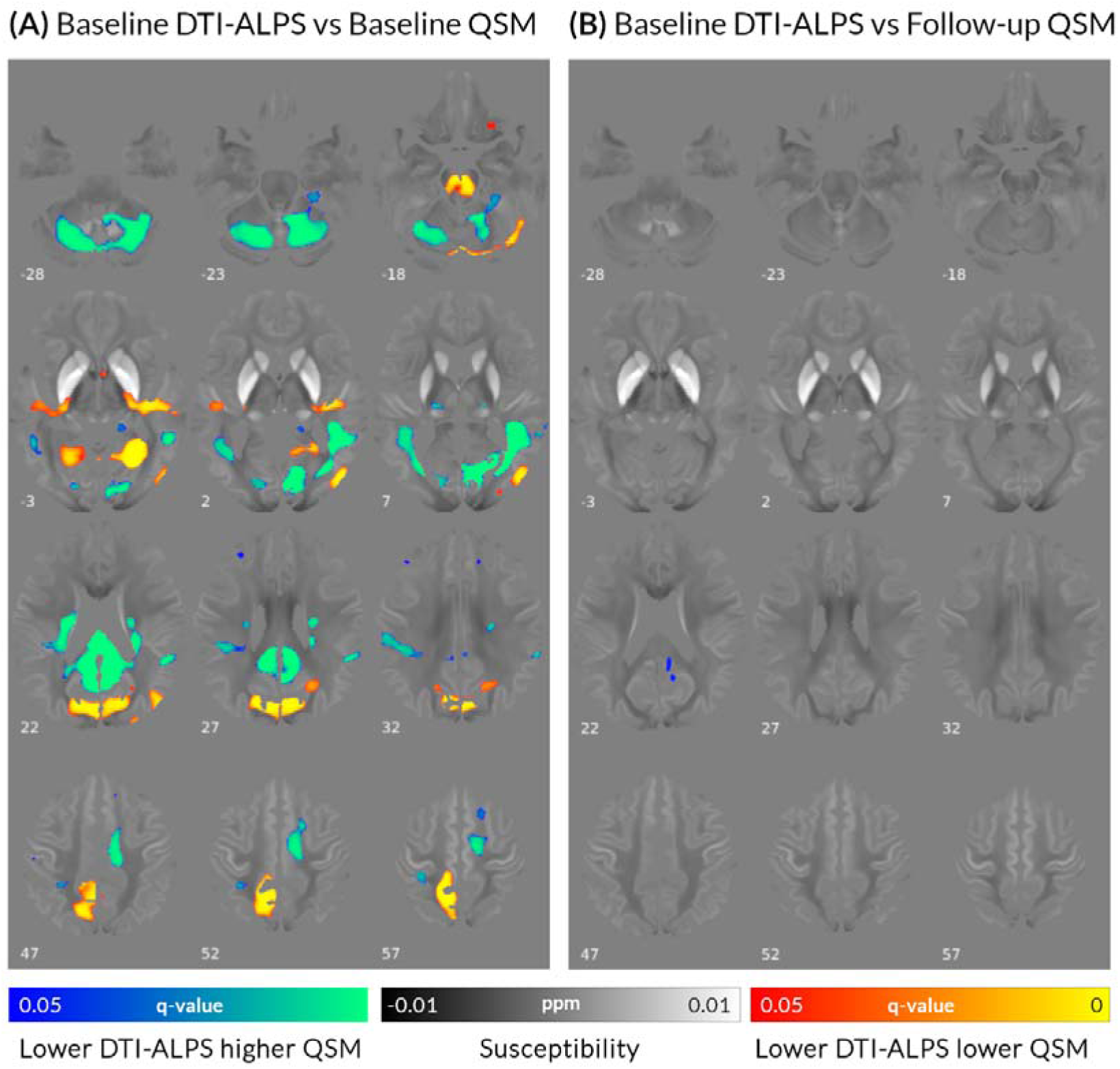
Lower DTI-ALPS index was associated with bidirectional changes in QSM levels in patients with Parkinson’s disease. **A. Baseline QSM.** In patients with Parkinson’s disease (n=98), lower baseline DTI-ALPS index was associated with both increases (blue colour scale) and reductions (red colour scale) in baseline susceptibility, adjusted for age and sex. **B. Follow-up increases in QSM.** Lower baseline DTI-ALPS index was associated increases in susceptibility after 3-year follow-up (Session 3) in patients with Parkinson’s disease (n=59), adjusted for age, sex and time between scans. Results are displayed on the QSM study-template in MNI152 space; numbers represent axial slice location in MNI152 space. Red/yellow clusters represent voxels where a significant positive correlation was seen with DTI-ALPS index (family-wise error corrected p-value, q<0.05) and Blue/green clusters voxels where a significant negative correlation was seen (q<0.05). DTI-ALPS: diffusion tensor image analysis along the perivascular space; QSM: quantitative susceptibility mapping.

### DTI-ALPS and cortical thickness

We assessed the relationship between DTI-ALPS index and whole-brain cortical thickness in PD patients at baseline (n=98) and longitudinally (n=57). At baseline, there were no statistically significant correlations. However, lower baseline DTI-ALPS was associated with increased longitudinal cortical thinning (annualised percentage reduction) in a cluster within the pre-central gyrus (*Figure 4*).

**Figure 4.**
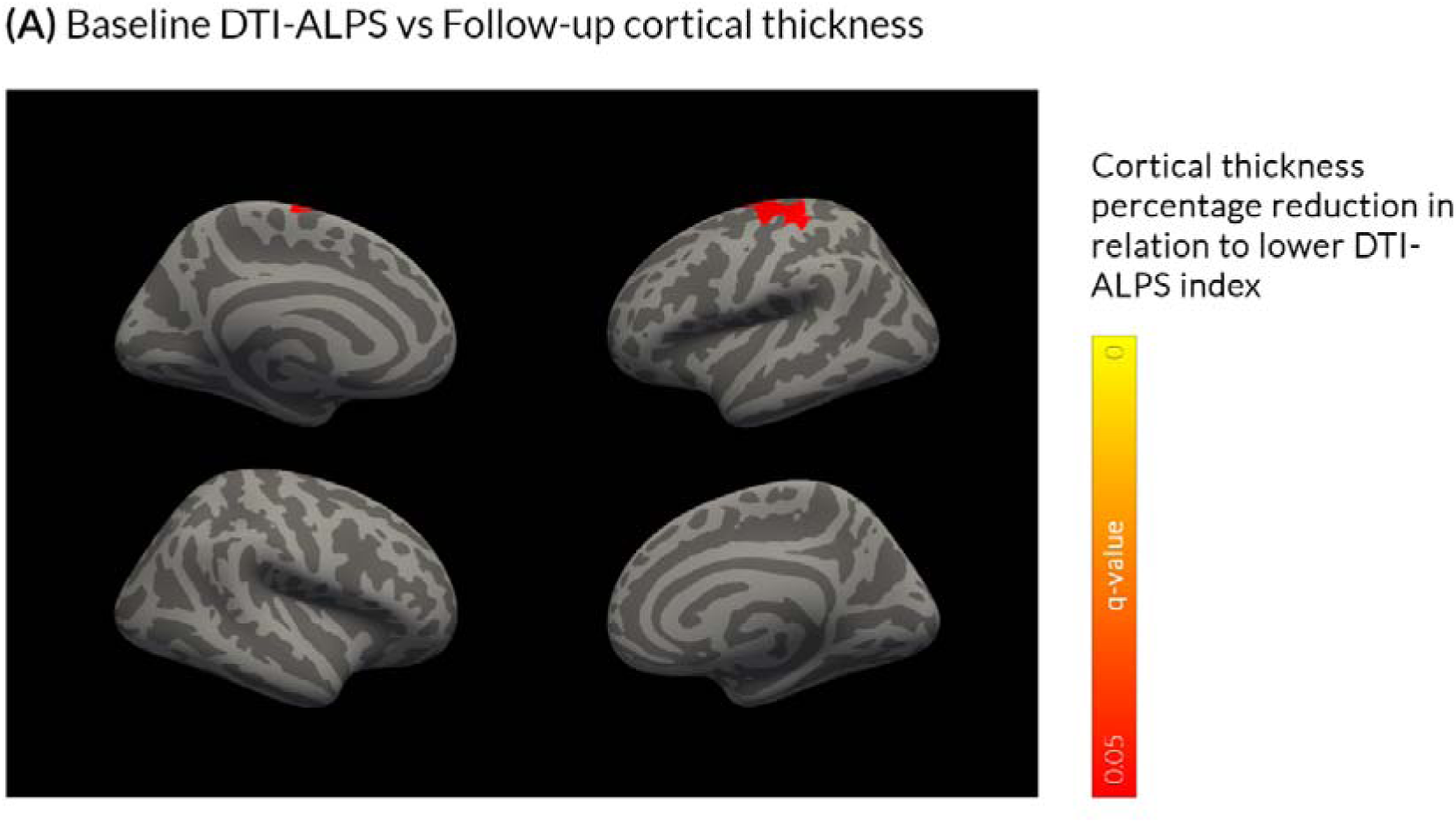
Lower DTI-ALPS index was associated with increased cortical thinning at follow-up. In patients with Parkinson’s disease, there was no correlation between a lower baseline DTI-ALPS index with baseline cortical thickness (whole brain analysis, family-wise error corrected p-value, q<0.05, age and sex corrected) but there was an association with percentage reduction in the pre-central gyrus after 3-year follow-up. DTI-ALPS: diffusion tensor image analysis along the perivascular space.

## Discussion

The glymphatic system plays a key role in clearance of pathological proteins and other waste products from the brain and has been postulated to contribute to PD symptoms^24,25^ and cognitive impairment^29–31^. However, how and when glymphatic clearance dysfunction interacts with other pathologies that contribute to PD dementia is not known. Here, we used an indirect, non-invasive correlate of glymphatic clearance, the DTI-ALPS index, to examine the relationship of glymphatic clearance dysfunction, with white matter degeneration, iron and amyloid accumulation and grey matter atrophy. We show that lower DTI-ALPS index: 1) is associated with cognitive performance and predicts poor outcomes in PD; 2) is associated with white matter macrostructural changes (reductions in fibre cross-section) and changes in magnetic susceptibility throughout the brain; but 3) its contribution to cognitive decline is distinct to those processes.

First, we show that DTI-ALPS index is reduced already at baseline in PD patients who progress to poor outcomes within 3 years, with additional reductions in those individuals during follow-up. In contrast, it remained relatively stable in PD patients with stable cognition and controls. The DTI-ALPS index was significantly associated with cognition and global disease burden in PD. This is in keeping with previous studies showing lower DTI-ALPS index with worsening global disease severity^24,59^ and cognition^25^ in PD. Other indirect methods of assessing glymphatic clearance also correlate with worse cognition in PD. Higher burden of enlarged perivascular spaces in the basal ganglia correlates to disease severity^24^ and cognitive performance^60^ in PD; and predicts cognitive decline over 5 years^61^. Glymphatic clearance relies on CSF movement through the perivascular space and enlargement of perivascular spaces is thought to reflect glymphatic dysfunction^62^. Polymorphisms of aquaporin-4, which play a crucial role in glymphatic function, are also linked to different rates of cognitive decline in PD^63^. Our study adds to the growing body of evidence that impaired glymphatic clearance is linked to cognitive impairment in PD.

Several hypotheses have been postulated on how impaired glymphatic clearance could contribute to neurodegeneration in PD, mainly focusing on the potential role of the glymphatic system in pathological protein aggregation^9,64^. Evidence from animal models suggests that suppression of glymphatic clearance leads to accumulation of both alpha-synuclein^13^ and beta-amyloid^15^, whilst misfolded protein spread via the glymphatic system has also been postulated^9,12^. In addition, lymphatic drainage seems to contribute to iron removal from the central nervous system in rats following intraventricular haemorrhage^65^, suggesting a possible role in iron clearance; and glymphatic dysfunction may trigger neuroinflammation^66^. All these processes may contribute in PD dementia: alpha-synuclein, beta-amyloid and tau accumulation, and cerebrovascular changes are seen in PD patients at autopsy^33^. Increased iron accumulation, particularly within the substantia nigra is also seen^67^ and may play a role in cell stress and atrophy^68^ and promoting alpha-synuclein aggregation^69^. Neuroinflammatory response contributes to dopaminergic cell loss and neurodegeneration^70^. These pathological processes lead to structural and functional brain changes and eventually, cognitive deficits in PD. However, few studies have tried to assess directly how impaired glymphatic clearance contributes to the structural changes seen in PD dementia. Here, we addressed this by assessing the relationship between DTI-ALPS index and established imaging markers that change in association with cognition in PD.

We found that lower baseline DTI-ALPS index, an indirect marker of poor glymphatic clearance, is associated with reductions in fibre cross-section, both at baseline and after 3-years. However, these changes were less diffuse and involved different tracts than those we have previously shown in PD with cognitive impairment using the same methodology^4,49^. Additionally, we found no relationship between DTI-ALPS and plasma NFL, despite the consistent observation of higher plasma NFL in association with cognitive decline in PD^4,71–73^. Another study in genetic frontotemporal dementia failed to find a correlation between plasma NFL and DTI-ALPS^74^. A potential explanation is different timing of glymphatic clearance and white matter degeneration. In Alzheimer’s, impaired amyloid clearance predates accumulation and overt neurodegeneration^75,76^; this could also be the case in PD. Glymphatic dysfunction correlates with degree of vascular damage to the white matter in health, as lower DTI-ALPS index and higher choroid plexus volume are associated with higher white matter hyperintensity volume at baseline and after 1-year^77^. In contrast, changes in fibre cross-section correlate with pathological protein distribution rather than white matter hyperintensities^51,78^, and NFL is a marker of axonal damage^79,80^ reflecting different mechanisms than white matter hyperintensities^81^. Therefore, another potential explanation for our findings is that DTI-ALPS predominantly affects white matter through cerebrovascular changes, which are not captured by fibre cross-section or NFL. This could be tested in future work specifically examining the relationship between the DTI-ALPS index and measures of small vessel disease in PD.

We found that lower DTI-ALPS index was associated with diffuse changes in QSM signal, with both areas of lower and higher magnetic susceptibility at baseline, and these also involved different regions than those previously linked to cognition in PD^6,42^. Only one prior study has concurrently assessed DTI-ALPS and magnetic susceptibility: Zhou et al. found a correlation between basal ganglia QSM and DTI-ALPS adjusting for age and sex, in 213 older adults^32^. This was interpreted as evidence that the glymphatic system plays a role in brain iron accumulation; however, that was a cross-sectional association in healthy individuals using region-of-interest analysis and did not assess whether these changes were behaviourally relevant. Our findings of bidirectional magnetic susceptibility changes in relation to the DTI-ALPS index, in keeping with human^32^ and animal evidence^65^, suggests that indeed glymphatic clearance plays a role in brain iron distribution. However, given the differential spatial distribution and lack of mediating effect of DTI-ALPS on QSM, our findings do not support the notion that impaired glymphatic clearance is a direct cause of the iron accumulation seen in PD with cognitive impairment.

Perhaps surprisingly, we did not find a correlation between DTI-ALPS index and plasma p-tau181. Impaired glymphatic clearance leads to beta-amyloid and tau accumulation in animal models^14,15^, and the DTI-ALPS index is linked to amyloid- and tau-PET positivity in Alzheimer’s^28,29^. However in PD, plasma p-tau181 levels do not correlate with cognitive severity, neither in our cohort, nor in others^4,72,73^. This could be due to higher heterogeneity of beta-amyloid and tau pathology in PD which may, in some cases, be incidental rather than driving cognitive decline. Alternatively other p-tau measurements, such as p-tau217, with increased sensitivity in detecting amyloid pathology^82^ may have greater ability to identify an association.

We found little evidence for an association between DTI-ALPS index and cortical thickness in PD: with no relationship at baseline and lower DTI-ALPS index predicting accelerated cortical thinning only within the precentral gyrus at follow-up. Previous studies have shown a correlation between DTI-ALPS index and cortical grey matter volume in Alzheimer’s^29,30^ and cortical thickness in frontotemporal dementia^83^. However, all studies used a region of interest, rather than whole-brain approach in conditions where cortical atrophy patterns are well-established and preserved across individuals. In contrast, cortical atrophy is highly heterogeneous in PD, occurring later in the disease course^26,84–87^.

Across all tested modalities, the spatial profiles of changes in relation to DTI-ALPS were different to those seen in relation to cognitive decline. This result, confirmed by the lack of mediation effect of DTI-ALPS to either fibre cross-section or QSM, suggests that glymphatic clearance does not drive white matter axonal degeneration nor brain iron accumulation in PD, but may play an independent role in cognitive impairment. An alternative explanation is that DTI-ALPS index is not a robust marker for glymphatic clearance. The DTI-ALPS index is deductive, and only indirectly reflects glymphatic clearance. It is affected by other factors that influence DWI signal^88,89^. Additionally, it can only be calculated within a specific region, at the level of the body of the lateral ventricles. Therefore it cannot assess regional differences in glymphatic function^88,89^. Finally, it has not been extensively validated in pathophysiological studies. Notwithstanding these limitations, the DTI-ALPS index is reduced under conditions where glymphatic function is impaired^21,31^ and it correlates strongly with intrathecal contrast administration^18^, the gold-standard in-vivo assessment method of glymphatic function. It is therefore likely to reflect glymphatic clearance, albeit not fully or solely. Other non-invasive methods to assess the glymphatic system are being developed, including perivascular spaces and arterial spin labelling^88,90^. Future work employing multiple metrics of glymphatic clearance could help elucidate its role in PD dementia.

Our study has some limitations. We did not acquire T2 or FLAIR imaging therefore could not quantify concurrent small vessel disease. Small vessel disease relates to cognition in PD^91,92^ and negatively correlates with DTI-ALPS^77,93^; it is possible that small vessel disease mediates DTI-ALPS’ effect on cognition in PD, and this could be examined in future work. Although ours was a longitudinal cohort, only group-level effects can be examined in our study design and whether DTI-ALPS is useful at the individual level remains to be determined. Additionally, this was a single-centre study and findings should be validated in additional populations.

### Conclusions

The DTI-ALPS index is lower in PD patients who progress to poor outcomes over 3-years and correlates with cognition. We show that although the DTI-ALPS index is linked to poor cognitive outcomes in PD, it is not the driver of the white matter axonal degeneration, or the iron accumulation seen in PD patients with cognitive impairment. Whether this reflects limitation of the technique or an independent role of impaired glymphatic clearance in Parkinson’s dementia remains to be seen.

## Acknowledgements

We thank our participants for their time and effort. We acknowledge the support of NVIDIA Corporation with the donation of the Quadro P6000 GPU used for this research. The authors acknowledge the use of the UCL Myriad High Performance Computing Facility (Myriad@UCL), and associated support services, in the completion of this work.

## References

1. Williams-Gray CH, Mason SL, Evans JR, et al. The CamPaIGN study of Parkinson’s disease: 10-year outlook in an incident population-based cohort. J. Neurol. Neurosurg. Psychiatry 2013;84(11):1258–1264.

2. Aarsland D, Brønnick K, Ehrt U, et al. Neuropsychiatric symptoms in patients with Parkinson’s disease and dementia: frequency, profile and associated care giver stress. J. Neurol. Neurosurg. Psychiatry 2007;78(1):36–42.

3. Dorsey ER, Elbaz A, Nichols E, et al. Global, regional, and national burden of Parkinson’s disease, 1990–2016: a systematic analysis for the Global Burden of Disease Study 2016. Lancet Neurol. 2018;17(11):939–953.

4. Zarkali A, Hannaway N, McColgan P, et al. Neuroimaging and plasma marker evidence for white matter macrostructure loss in Parkinson’s disease [Internet]. 2024;fcae130[cited 2023 Oct 18] Available from: https://www.biorxiv.org/content/10.1101/2023.09.22.558937v1

5. Cheng HC, Ulane CM, Burke RE. Clinical progression in parkinson disease and the neurobiology of axons. Ann. Neurol. 2010;67(6):715–725.

6. Thomas GEC, Leyland LA, Schrag A-E, et al. Brain iron deposition is linked with cognitive severity in Parkinson’s disease. J. Neurol. Neurosurg. Psychiatry 2020;91(4):418–425.

7. Baik K, Kim HR, Park M, et al. Effect of amyloid on cognitive performance in parkinson’s disease and dementia with lewy bodies. Mov. Disord. Off. J. Mov. Disord. Soc. 2023;38(2):278–285.

8. Hijazi Z, Yassi N, O’Brien JT, Watson R. The influence of cerebrovascular disease in dementia with Lewy bodies and Parkinson’s disease dementia. Eur. J. Neurol. 2022;29(4):1254–1265.

9. Nedergaard M, Goldman SA. Glymphatic failure as a final common pathway to dementia. Science 2020;370(6512):50–56.

10. Iliff JJ, Wang M, Liao Y, et al. A Paravascular Pathway Facilitates CSF Flow Through the Brain Parenchyma and the Clearance of Interstitial Solutes, Including Amyloid β Sci. Transl. Med. 2012;4(147):147ra111–147ra111.

11. Eide PK, Ringstad G. Functional analysis of the human perivascular subarachnoid space. Nat. Commun. 2024;15(1):2001.

12. Fujita K, Homma H, Jin M, et al. Mutant α synuclein propagates via the lymphatic system of the brain in the monomeric state [Internet]. Cell Rep. 2023;42(8)[cited 2024 Sep 30] Available from: https://www.cell.com/cell-reports/abstract/S2211-1247(23)00973-7

13. Zhang Y, Zhang C, He X-Z, et al. Interaction Between the Glymphatic System and α Synuclein in Parkinson’s Disease. Mol. Neurobiol. 2023;60(4):2209–2222.

14. Harrison IF, Ismail O, Machhada A, et al. Impaired glymphatic function and clearance of tau in an Alzheimer’s disease model. Brain J. Neurol. 2020;143(8):2576–2593.

15. Xu Z, Xiao N, Chen Y, et al. Deletion of aquaporin-4 in APP/PS1 mice exacerbates brain Aβ accumulation and memory deficits. Mol. Neurodegener. 2015;10:58.

16. Bohr T, Hjorth PG, Holst SC, et al. The glymphatic system: Current understanding and modeling. iScience 2022;25(9):104987.

17. Taoka T, Masutani Y, Kawai H, et al. Evaluation of glymphatic system activity with the diffusion MR technique: diffusion tensor image analysis along the perivascular space (DTI-ALPS) in Alzheimer’s disease cases. Jpn. J. Radiol. 2017;35(4):172–178.

18. Zhang W, Zhou Y, Wang J, et al. Glymphatic clearance function in patients with cerebral small vessel disease. NeuroImage 2021;238:118257.

19. Tian Y, Cai X, Zhou Y, et al. Impaired glymphatic system as evidenced by low diffusivity along perivascular spaces is associated with cerebral small vessel disease: a population-based study. Stroke Vasc. Neurol. 2023;8(5):e002191.

20. Liu X, Barisano G, Shao X, et al. Cross-Vendor Test-Retest Validation of Diffusion Tensor Image Analysis along the Perivascular Space (DTI-ALPS) for Evaluating Glymphatic System Function. Aging Dis. 2024;15(4):1885–1898.

21. Clark O, Delgado-Sanchez A, Cullell N, et al. Diffusion tensor imaging analysis along the perivascular space in the UK biobank. Sleep Med. 2024;119:399–405.

22. Ma X, Li S, Li C, et al. Diffusion Tensor Imaging Along the Perivascular Space Index in Different Stages of Parkinson’s Disease [Internet]. Front. Aging Neurosci. 2021;13[cited 2024 Oct 4] Available from: https://www.frontiersin.org/journals/aging-neuroscience/articles/10.3389/fnagi.2021.773951/full

23. Si X, Guo T, Wang Z, et al. Neuroimaging evidence of glymphatic system dysfunction in possible REM sleep behavior disorder and Parkinson’s disease. NPJ Park. Dis. 2022;8(1):54.

24. Meng J-C, Shen M-Q, Lu Y-L, et al. Correlation of glymphatic system abnormalities with Parkinson’s disease progression: a clinical study based on non-invasive fMRI. J. Neurol. 2024;271(1):457–471.

25. He P, Shi L, Li Y, et al. The Association of the Glymphatic Function with Parkinson’s Disease Symptoms: Neuroimaging Evidence from Longitudinal and Cross-Sectional Studies. Ann. Neurol. 2023;94(4):672–683.

26. Zarkali A, Thomas GEC, Zetterberg H, Weil RS. Neuroimaging and fluid biomarkers in Parkinson’s disease in an era of targeted interventions. Nat. Commun. 2024;15(1):5661.

27. Kamagata K, Andica C, Takabayashi K, et al. Association of MRI Indices of Glymphatic System With Amyloid Deposition and Cognition in Mild Cognitive Impairment and Alzheimer Disease. Neurology 2022;99(24):e2648–e2660.

28. Hsu J-L, Wei Y-C, Toh CH, et al. Magnetic Resonance Images Implicate That Glymphatic Alterations Mediate Cognitive Dysfunction in Alzheimer Disease. Ann. Neurol. 2023;93(1):164–174.

29. Huang S-Y, Zhang Y-R, Guo Y, et al. Glymphatic system dysfunction predicts amyloid deposition, neurodegeneration, and clinical progression in Alzheimer’s disease. Alzheimers Dement. J. Alzheimers Assoc. 2024;20(5):3251–3269.

30. Chang H-I, Huang C-W, Hsu S-W, et al. Gray matter reserve determines glymphatic system function in young-onset Alzheimer’s disease: Evidenced by DTI-ALPS and compared with age-matched controls. Psychiatry Clin. Neurosci. 2023;77(7):401–409.

31. Siow TY, Toh CH, Hsu J-L, et al. Association of Sleep, Neuropsychological Performance, and Gray Matter Volume With Glymphatic Function in Community-Dwelling Older Adults. Neurology 2022;98(8):e829–e838.

32. Zhou W, Shen B, Shen W, et al. Dysfunction of the Glymphatic System Might Be Related to Iron Deposition in the Normal Aging Brain. Front. Aging Neurosci. 2020;12:559603.

33. Irwin DJ, Grossman M, Weintraub D, et al. Neuropathological and genetic correlates of survival and dementia onset in synucleinopathies: a retrospective analysis. Lancet Neurol. 2017;16(1):55–65.

34. Hannaway N, Zarkali A, Leyland L-A, et al. Visual dysfunction is a better predictor than retinal thickness for dementia in Parkinson’s disease [Internet]. J. Neurol. Neurosurg. Psychiatry 2023;[cited 2023 Jul 14] Available from: https://jnnp.bmj.com/content/early/2023/04/20/jnnp-2023-331083

35. Postuma RB, Berg D, Stern M, et al. MDS clinical diagnostic criteria for Parkinson’s disease. Mov. Disord. 2015;30(12):1591–1601.

36. Creavin ST, Wisniewski S, Noel-Storr AH, et al. Mini-Mental State Examination (MMSE) for the detection of dementia in clinically unevaluated people aged 65 and over in community and primary care populations [Internet]. Cochrane Database Syst. Rev. 2016;(1)[cited 2019 Feb 15] Available from: https://www.cochranelibrary.com/cdsr/doi/10.1002/14651858.CD011145.pub2/abstract

37. Dalrymple-Alford JC, MacAskill MR, Nakas CT, et al. The MoCA: well-suited screen for cognitive impairment in Parkinson disease. Neurology 2010;75(19):1717–1725.

38. Goetz CG, Tilley BC, Shaftman SR, et al. Movement Disorder Society-sponsored revision of the Unified Parkinson’s Disease Rating Scale (MDS-UPDRS): scale presentation and clinimetric testing results. Mov. Disord. Off. J. Mov. Disord. Soc. 2008;23(15):2129–2170.

39. Shumway-Cook A, Brauer S, Woollacott M. Predicting the Probability for Falls in Community-Dwelling Older Adults Using the Timed Up & Go Test. Phys. Ther. 2000;80(9):896–903.

40. Tomlinson CL, Stowe R, Patel S, et al. Systematic review of levodopa dose equivalency reporting in Parkinson’s disease. Mov. Disord. 2010;25(15):2649–2653.

41. Litvan I, Goldman JG, Tröster AI, et al. Diagnostic Criteria for Mild Cognitive Impairment in Parkinson’s Disease: Movement Disorder Society Task Force Guidelines. Mov. Disord. Off. J. Mov. Disord. Soc. 2012;27(3):349–356.

42. Thomas GEC, Hannaway N, Zarkali A, et al. Longitudinal Associations of Magnetic Susceptibility with Clinical Severity in Parkinson’s Disease [Internet]. Mov. Disord. [date unknown];n/a(n/a)[cited 2024 Mar 4] Available from: https://onlinelibrary.wiley.com/doi/abs/10.1002/mds.29702

43. Veraart J, Fieremans E, Novikov DS. Diffusion MRI noise mapping using random matrix theory. Magn. Reson. Med. 2016;76(5):1582–1593.

44. Kellner E, Dhital B, Kiselev VG, Reisert M. Gibbs-ringing artifact removal based on local subvoxel-shifts. Magn. Reson. Med. 2016;76(5):1574–1581.

45. Anderson G. Assuring Quality/Resisting Quality Assurance: Academics’ responses to ‘quality’ in some Australian universities. Qual. High. Educ. 2006;12(2):161–173.

46. Tustison NJ, Avants BB, Cook PA, et al. N4ITK: improved N3 bias correction. IEEE Trans. Med. Imaging 2010;29(6):1310–1320.

47. Hollander, T., Raffelt D, Connelly A. Unsupervised 3-tissue response function estimation from single-shell or multi-shell diffusion MR data without a co-registered T1 image. ISMRM Workshop Break. Barriers Diffus. MRI 2016;5.

48. Raffelt D, Tournier J-D, Fripp J, et al. Symmetric diffeomorphic registration of fibre orientation distributions. NeuroImage 2011;56(3):1171–1180.

49. Zarkali A, McColgan P, Leyland L-A, et al. Visual Dysfunction Predicts Cognitive Impairment and White Matter Degeneration in Parkinson’s Disease [Internet]. Mov. Disord. 2021;n/a(n/a)[cited 2021 Jan 22] Available from: https://movementdisorders.onlinelibrary.wiley.com/doi/abs/10.1002/mds.28477

50. Rau Y-A, Wang S-M, Tournier J-D, et al. A longitudinal fixel-based analysis of white matter alterations in patients with Parkinson’s disease [Internet]. NeuroImage Clin. 2019;24[cited 2020 Jul 17] Available from: https://www.ncbi.nlm.nih.gov/pmc/articles/PMC6889638/

51. Dewenter A, Jacob MA, Cai M, et al. Disentangling the effects of Alzheimer’s and small vessel disease on white matter fibre tracts. Brain 2022;146(2):678–689.

52. Reuter M, Schmansky NJ, Rosas HD, Fischl B. Within-subject template estimation for unbiased longitudinal image analysis. NeuroImage 2012;61(4):1402–1418.

53. Dymerska B, Eckstein K, Bachrata B, et al. Phase unwrapping with a rapid opensource minimum spanning tree algorithm (ROMEO). Magn. Reson. Med. 2021;85(4):2294– 2308.

54. Smith SM. Fast robust automated brain extraction. Hum. Brain Mapp. 2002;17(3):143– 155.

55. Zhou D, Liu T, Spincemaille P, Wang Y. Background field removal by solving the Laplacian boundary value problem. NMR Biomed. 2014;27(3):312–319.

56. Acosta-Cabronero J, Milovic C, Mattern H, et al. A robust multi-scale approach to quantitative susceptibility mapping. NeuroImage 2018;183:7–24.

57. Smith RE, Tournier J-D, Calamante F, Connelly A. SIFT: Spherical-deconvolution informed filtering of tractograms. NeuroImage 2013;67:298–312.

58. Mori S, Wakana S, van Zijl P, Nagae-Poetscher L. MRI Atlas of Human White Matter - 1st Edition [Internet]. Amsterdam: Elsevier; [date unknown].[cited 2019 May 9] Available from: https://www.elsevier.com/books/mri-atlas-of-human-white-matter/mori/978-0-444-51741-8

59. Ma X, Li S, Li C, et al. Diffusion Tensor Imaging Along the Perivascular Space Index in Different Stages of Parkinson’s Disease. Front. Aging Neurosci. 2021;13:773951.

60. Donahue EK, Foreman RP, Duran JJ, et al. Increased perivascular space volume in white matter and basal ganglia is associated with cognition in Parkinson’s Disease. Brain Imaging Behav. 2024;18(1):57–65.

61. Park YW, Shin N-Y, Chung SJ, et al. Magnetic Resonance Imaging-Visible Perivascular Spaces in Basal Ganglia Predict Cognitive Decline in Parkinson’s Disease. Mov. Disord. Off. J. Mov. Disord. Soc. 2019;34(11):1672–1679.

62. Weller RO, Hawkes CA, Kalaria RN, et al. White Matter Changes in Dementia: Role of Impaired Drainage of Interstitial Fluid. Brain Pathol. 2015;25(1):63–78.

63. Fang Y, Dai S, Jin C, et al. Aquaporin-4 Polymorphisms Are Associated With Cognitive Performance in Parkinson’s Disease. Front. Aging Neurosci. 2021;13:740491.

64. Ryman SG, Vakhtin AA, Mayer AR, et al. Abnormal Cerebrovascular Activity, Perfusion, and Glymphatic Clearance in Lewy Body Diseases. Mov. Disord. 2024;39(8):1258– 1268.

65. Bocheng W, Chaofeng L, Chuan C, et al. Intracranial lymphatic drainage discharges iron from the ventricles and reduce the occurrence of chronic hydrocephalus after intraventricular hemorrhage in rats. Int. J. Neurosci. 2020;130(2):130–135.

66. Ryman SG, Vakhtin AA, Mayer AR, et al. Abnormal Cerebrovascular Activity, Perfusion, and Glymphatic Clearance in Lewy Body Diseases. Mov. Disord. 2024;39(8):1258– 1268.

67. Langkammer C, Schweser F, Krebs N, et al. Quantitative susceptibility mapping (QSM) as a means to measure brain iron? A post mortem validation study. NeuroImage 2012;62(3):1593–1599.

68. Li J, Cao F, Yin H, et al. Ferroptosis: past, present and future. Cell Death Dis. 2020;11(2):88–88.

69. Ostrerova-Golts N, Petrucelli L, Hardy J, et al. The A53T alpha-synuclein mutation increases iron-dependent aggregation and toxicity. J. Neurosci. 2000;20(16):6048–54.

70. Q W, J Z, S P, et al. The link between neuroinflammation and the neurovascular unit in synucleinopathies [Internet]. Sci. Adv. 2023;9(7)[cited 2024 Oct 10] Available from: https://pubmed.ncbi.nlm.nih.gov/36791205/

71. Aamodt WW, Waligorska T, Shen J, et al. Neurofilament Light Chain as a Biomarker for Cognitive Decline in Parkinson Disease. Mov. Disord. Off. J. Mov. Disord. Soc. 2021;36(12):2945–2950.

72. Batzu L, Rota S, Hye A, et al. Plasma p-tau181, neurofilament light chain and association with cognition in Parkinson’s disease. Npj Park. Dis. 2022;8(1):1–7.

73. Pagonabarraga J, Pérez-González R, Bejr-kasem H, et al. Dissociable contribution of plasma NfL and p-tau181 to cognitive impairment in Parkinson’s disease. Parkinsonism Relat. Disord. 2022;105:132–138.

74. Premi E, Diano M, Mattioli I, et al. Impaired glymphatic system in genetic frontotemporal dementia: a GENFI study. Brain Commun. 2024;6(4):fcae185.

75. Mawuenyega KG, Sigurdson W, Ovod V, et al. Decreased clearance of CNS beta-amyloid in Alzheimer’s disease. Science 2010;330(6012):1774.

76. Patterson BW, Elbert DL, Mawuenyega KG, et al. Age and amyloid effects on human central nervous system amyloid-beta kinetics. Ann. Neurol. 2015;78(3):439–453.

77. Y L, Y Z, W Z, et al. Choroid Plexus Enlargement Exacerbates White Matter Hyperintensity Growth through Glymphatic Impairment [Internet]. Ann. Neurol. 2023;94(1)[cited 2024 Oct 9] Available from: https://pubmed.ncbi.nlm.nih.gov/36971336/

78. Ahmadi K, Pereira JB, Westen D van, et al. Fixel-based analysis reveals macrostructural white matter changes associated with tau pathology in early stages of Alzheimer’s disease [Internet]. 2023;2023.02.17.23286094.[cited 2023 Nov 15] Available from: https://www.medrxiv.org/content/10.1101/2023.02.17.23286094v1

79. Zetterberg H, Skillbäck T, Mattsson N, et al. Association of Cerebrospinal Fluid Neurofilament Light Concentration With Alzheimer Disease Progression. JAMA Neurol. 2016;73(1):60–67.

80. Sjögren M, Blomberg M, Jonsson M, et al. Neurofilament protein in cerebrospinal fluid: a marker of white matter changes. J. Neurosci. Res. 2001;66(3):510–516.

81. Dhana A, DeCarli C, Dhana K, et al. White matter hyperintensity, neurofilament light chain, and cognitive decline. Ann. Clin. Transl. Neurol. 2022;10(3):321–327.

82. Kivisäkk P, Fatima HA, Cahoon DS, et al. Clinical evaluation of a novel plasma pTau217 electrochemiluminescence immunoassay in Alzheimer’s disease. Sci. Rep. 2024;14(1):629.

83. Xiao D, Li J, Ren Z, et al. Association of cortical morphology, white matter hyperintensity, and glymphatic function in frontotemporal dementia variants. Alzheimers Dement. 2024;20(9):6045–6059.

84. Zarkali A, Adams RA, Psarras S, et al. Increased weighting on prior knowledge in Lewy body-associated visual hallucinations [Internet]. Brain Commun. 2019;1(1)[cited 2019 Oct 29] Available from: https://academic.oup.com/braincomms/article/1/1/fcz007/5532510

85. Marques A, Macias E, Pereira B, et al. Volumetric changes and clinical trajectories in Parkinson’s disease: a prospective multicentric study [Internet]. J. Neurol. 2023;[cited 2023 Oct 9] Available from: 10.1007/s00415-023-11947-0

86. Park C, Shin N-Y, Yoo S-W, et al. Simulating the progression of brain structural alterations in Parkinson’s disease. Npj Park. Dis. 2022;8(1):1–8.

87. Pietracupa S, Belvisi D, Piervincenzi C, et al. White and gray matter alterations in de novo PD patients: which matter most? J. Neurol. 2023;270(5):2734–2742.

88. Taoka T, Ito R, Nakamichi R, et al. Diffusion Tensor Image Analysis ALong the Perivascular Space (DTI-ALPS): Revisiting the Meaning and Significance of the Method. Magn. Reson. Med. Sci. 2024;23(3):268–290.

89. Ringstad G. Glymphatic imaging: a critical look at the DTI-ALPS index. Neuroradiology 2024;66(2):157–160.

90. Perera C, Cruz R, Shemesh N, et al. Non-invasive MRI of Blood-Cerebrospinal Fluid-Barrier Function: a Functional Biomarker of Early Alzheimer’s Disease Pathology [Internet]. 2024;2024.03.06.583668.[cited 2024 Oct 10] Available from: https://www.biorxiv.org/content/10.1101/2024.03.06.583668v1

91. Kummer BR, Diaz I, Wu X, et al. Associations between Cerebrovascular Risk Factors and Parkinson Disease. Ann. Neurol. 2019;86(4):572–581.

92. Toledo JB, Arnold SE, Raible K, et al. Contribution of cerebrovascular disease in autopsy confirmed neurodegenerative disease cases in the National Alzheimer’s Coordinating Centre. Brain 2013;136(9):2697–2706.

93. Zhang W, Zhou Y, Wang J, et al. Glymphatic clearance function in patients with cerebral small vessel disease. NeuroImage 2021;238:118257.

