## Supplementary Material for "Impaired glymphatic clearance independently contributes to poor outcomes in Parkinson’s disease"

### **S1. Mediation analyses between glymphatic function (DTI-ALPS index), white matter degeneration (fibre cross-section), grey matter atrophy (cortical thickness), iron accumulation (QSM) and cognitive outcomes in Parkinson’s disease**

**Methods**

To explore the interplay between glymphatic function, white matter degeneration, iron accumulation and poor cognitive outcomes in Parkinson’s, we conducted a mediation analysis to derive total, direct and indirect effects. To perform mediation analysis, mean fibre cross-section, grey matter volume and quantitative susceptibility mapping (QSM) values were derived from regions of interest (ROIs) that were most related to cognition based on our previous work (Thomas et al., 2020; Zarkali et al., 2024, 2021): the corpus callosum for white matter, hippocampus for grey matter and nucleus basalis of Meynert (NBM) for QSM. Mean fibre cross-section was derived from the entirety of the corpus callosum using a mask from the JHU white matter atlas, mean bilateral hippocampal volume was derived from the built-in Freesurfer segmentation and mean signed QSM was derived from a manually traced mask of the NBM as previously described (Thomas et al., 2020).

We first assessed the inter-relationship between QSM, fibre cross-section, hippocampal volume and DTI-ALPS using partial correlation with age and sex as covariates as well as each variable’s relationship with change in combined cognitive scores (Session 3 – Baseline) and poor outcomes to select variables for mediation analysis. Only variables that were correlated with each other and the outcome of interest were included.

Mediation analysis was performed between change in 1) combined cognitive scores (Session 3 – Baseline) and baseline mean QSM and DTI-ALPS, with age and sex as covariates using linear regression and 2) poor outcomes and hippocampal volume and corpus callosum fibre cross-section using logistic regression.

**Results**

Mean corpus callosum fibre cross-section had a significant direct effect on poor outcomes (β=1.348, p=0.01) and a small effect on cortical thickness (β=0.001, p<0.001) (*Figure A*). QSM signal of the NBM had a direct effect on change in cognitive scores (β=-4.445, p=0.001). Neither QSM nor DTI-ALPS acted as a mediator for the other, suggesting independent contributions to
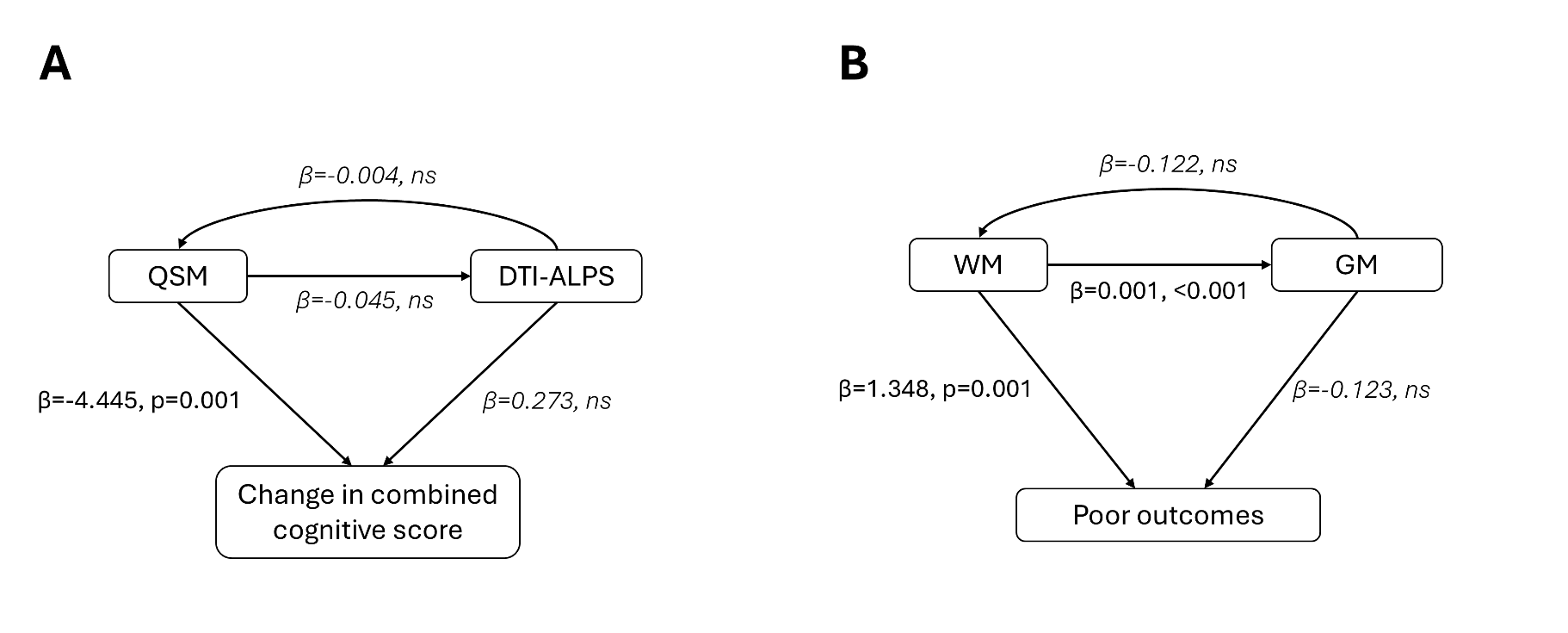
cognitive decline (*Figure B*).

**Figure. Mediation analysis of factors contributing to cognitive change in Parkinson’s.**

Standardized coefficients (β) were calculated for each association with (**A**) change in combined cognitive scores (Session 3 – Baseline) and (**B**) poor outcomes in patients with Parkinson’s disease, using mediation analysis of baseline values for each variable adjusted for age and sex.

DTI-ALPS: diffusion tensor image analysis along the perivascular space; GM: bilateral hippocampal volume; ns: non-significant (p-value>0.05); QSM: mean quantitative susceptibility mapping value of the nucleus basalis of Meynert; WM: mean fibre cross-section of the corpus callosum.
